# Recognition and identification of species in the *Bombus lucorum*-complex – A review and outlook

**DOI:** 10.1101/011379

**Authors:** Silas Bossert

## Abstract

The recognition of cryptic species represents one of the major challenges in current taxonomy and affects our understanding of global diversity. In practice, the process from discovery to acceptance in the scientific community can take an extensive length of time. A prime example is the traditionally difficult taxonomy of the cryptic bumblebee species belonging to the *Bombus lucorum*-complex. The status of the three European species in the group – *Bombus lucorum* and the closely related *Bombus cryptarum* and *Bombus magnus* – has recently become widely accepted, primarily due to investigations of nucleotide sequences and marking pheromones. In contrast, doubts prevail concerning the validity of species identification based on morphology. As a consequence, our knowledge of the species is muddled in a mire of unreliable and confusing literature data from a large number of authors over the centuries. To clarify this issue, this paper provides a recapitulation of the historical literature and highlights the milestones in the process of species recognition. Further, the possibility of a morphologically based species identification is discussed in the context of new molecular data. Finally, this review outlines the current challenges and provides directions for future issues.

## Introduction

Bumblebees (*Bombus*, Latreille 1802) are considered to be a striking feature of Europe’s pollinator fauna (e.g. Corbet et al. 1991, Neumayer and Paulus 1999, Goulson et al. 2007, Goulson 2010). In contrast to most other bee genera, bumblebees are referred to be intuitively recognizable and are rarely confused with other bees (Amiet 1996, Gokcezade et al. 2010, Amiet and Krebs 2012). Yet species determination requires expertise, and reliable identification in the field is often impossible. Reasons for this are the relatively monotonous morphology (Michener 2007), enormous variability in coloration and size which is often associated with biogeographical distribution (e.g. Vogt 1909, Vogt 1911, Krüger 1951, Løken 1973, Pekkarinen 1979) and the fact that the same or similar color-patterns are often repeated in various species (Dalla Torre 1880, Reinig 1939, Amiet 1996, Williams 2007 and references therein). One of these groups with very similar morphology consists of the European species of the subgenus *Bombus* s. str.: *Bombus terrestris* L., 1758, *B. lucorum* L., 1761, *B. cryptarum* Fabricius, 1775, *B. magnus* Vogt, 1911 and *B. sporadicus* Nylander, 1848. Two species of this group, *B. lucorum* and particularly *B. terrestris,* are of great economic interest since the extensive use of bumblebees for commercial greenhouse pollination (Velthuis and van Doorn 2006, Winter et al. 2006). In the past decades, there has been much disagreement on the taxonomy of this group. Especially the status of *B. lucorum* and the closely related *B. cryptarum* and *B. magnus*, forming the so-called *Bombus lucorum*-complex, has been intensively discussed. This can be traced to an exceptionally high degree of synonymisation: Williams (1998) reported far more than 100 infrasubspecific names just for *Bombus lucorum* s.l. In contrast, the species status of the three distinct species in Europe is widely accepted nowadays, primarily based on investigations of nucleotide sequences of the mitochondrial COI gene (Bertsch et al. 2005, Murray et al. 2008, Bertsch 2009, Carolan et al. 2012, Williams et al. 2012) and male labial gland secretions (Bertsch 1997, Bertsch et al. 2004, Bertsch et al. 2005). Still, serious doubts remain concerning the validity of species identification based on morphology and the reliability of certain distinguishing characters have been challenged (e.g. Williams 2000, Carolan et al. 2012). As a consequence of this doubtful delineation, our current knowledge about the species is muddled in a mire of unreliable literature data from numerous authors over the centuries. Only few studies on the species exist that are backed up by reliable species identification using biochemical methods. In addition, information about diagnostic characters in the literature are often confusing or based on insufficient underlying data sources. To rectify the problem, this review provides an overview on the species recognition and the distinguishability of the *Bombus lucorum*-complex. Further, it provides an urgently required reappraisal to pave the way for future investigations.

### Bombus lucorum vs. Bombus magnus

*Bombus lucorum and B. terrestris* were described by Linnaeus in 1761 and 1758, respectively. Their species status has been widely accepted in the last century. Only single authors doubted their status and lumped them together (e.g. Warncke 1981, Warncke 1986). More than a century later, *B. magnus* was described by Vogt (1911) in a single sentence as a ‘forma nova *magnus*’ without detailed information. It was probably the same species that was described as *Bombus terrestris* var. *flavoscutellaris* by Trautmann and Trautmann (1915). The species description of *B. magnus* was conducted by Krüger (1951, 1954) with detailed descriptions of all castes and several *races* and *ethna*, which are difficult to comprehend from today’s view. Some earlier experts failed to distinguish *B. lucorum* and *B. magnus* (Elfving 1960, Ander 1965), others primarily highlighted the need of further studies (Alford 1975, Delmas 1981). Løken (1973) conducted a grand morphometric analysis and advocated their species status, primarily based on measurements of queens, whereas the distinguishability of workers and males remained uncertain. Her work was confirmed and enhanced by further specific indices by Tkalců (1974). At that time the first biochemical results in form of male labial gland marking pheromones emerged (Kullenberg et al. 1970, Bergström et al. 1973, Bergström et al. 1981). For *B. lucorum,* two similar but distinctly different profiles could be identified concerning a ‘dark’ and a ‘blonde’ form, supporting Løken´s (1973) view. However, common to all of the above mentioned literature is the fact that a previously unknown species, *B. cryptarum*, occurs sympatrically with *B. lucorum* and *B. magnus* and probably biased their results due to a species mix in their samples. This is likely the reason why the results from Pekkarinen (1979) are not in line with the others. Even though other authors also overlooked a possible third taxon (Scholl and Obrecht 1983, Pamilo et al. 1984), their results based on enzyme electrophoresis were convincing that *B. lucorum* is not a single species.

### A third species comes into play

Using morphological and morphometric methods, Rasmont (1981a, 1981b) was the first who recognized a putative third species and described it as *Bombus lucocryptarum* Ball which was later synonymized with *Bombus cryptarum* Fabricius (Rasmont 1983). Interestingly this taxon was also previously described as *Bombus lucorum* var. *pseudocryptarum* Skorikov from Russia and Poland (Skorikov 1913). Rasmont (1981b) provided a determination table for the queens. Tables for both female castes (Rasmont 1984) and males (Rasmont et al. 1986) followed, even if those for the latter were of limited applicability. His keys were supported by remarkable crossing experiments between the three putative species (De Jonghe 1982, De Jonghe and Rasmont 1983, Rasmont and De Jonghe 1985). His cross breeding of the three putative taxa failed, even though copulation and egg deposition were observed. In contrast, his breeding within the examined taxa succeeded. In contrast, no interspecific mating was observed in the experiments of Bučánková et al. (2011). However, due to the limited sample size of the studies, the results strongly indicated reproductive isolation but are not completely reliable. In general, although the conviction that *B. lucorum* consisted of more than one taxon grew, the species were still lumped together by some authors (Warncke 1986, Westrich 1990). Williams (1991, 1998) provisionally synonymized the potential species. The confirmation of a third species with biochemical methods remained open for some time (Obrecht and Scholl 1984, Scholl et al. 1990, Scholl et al. 1992, Pamilo et al. 1997), probably due to the similar enzyme genetic profiles of *B. cryptarum* and *B. magnus*. However, it is likely that the samples of *B. cryptarum* and *B. magnus* used for analyses were mixed, a point that Bertsch et al. (2004) presupposed for Pamilo et al. (1997). With recurring theme, the morphometric attempts of Baker (1996) were of restricted value, since *B. cryptarum* was not considered as a separate species and the same applies for Macdonald (1999). He advocated *B. lucorum* and *B. magnus* as good species based on the coloration of the pile (extended yellow collar of queens of *B. magnus*; for a review of morphological traits see below) and observations concerning their ecology. In retrospect, it seems highly likely that at least some of the examined specimens from his study were in fact *B. cryptarum,* since this species occurs most frequently in the mainland of northern Scotland (Macdonald, personal communication). This is an explanation why Williams (2000) could not find a clear gap in collar extending between *B. lucorum* and *B. magnus*: *B. cryptarum* queens have on average a collar extension between the latter two species (Carolan et al. 2012) which may have critically biased the measurements. However, the first sufficient biochemical evidence for all three species was conducted by Bertsch (1997) and Bertsch et al. (2004) by the identification of three distinct male labial gland secretion profiles: the profiles of *B. cryptarum* and *B. magnus* are similar and share ethyl dodecanoate as the main component. Yet they clearly differ in minor components such as alcohols (Bertsch et al. 2004, Bertsch et al. 2005). Recently the great stability of the labial gland secretion composition of *B. cryptarum* over great geographical ranges was shown, a fact that supports their value for species recognition (Bertsch and Schweer 2012).

### Nucleotide sequence data improved the understanding

The debate gained new life with the application of phylogenetic analyses using nucleotide sequences of the mitochondrial cytochrome oxidase I gene (COI). With this method, the composition of three distinct molecular operational taxonomic units (MOTUs) in the European *B. lucorum*-complex was convincingly confirmed multiple times (Bertsch et al. 2005, Murray et al. 2008, Bertsch 2009, Carolan et al. 2012), even if the taxonomic state of knowledge remains incomplete in the global context (Williams et al. 2012). Admittedly, COI barcoding and its applicability for species recognition has been criticized (e.g. Will and Rubinoff 2004, DeSalle et al. 2005, Meyer and Paulay 2005, for a review see Taylor and Harris 2012), the results for this species complex seem convincing. The interspecific genetic divergences of the species are considerably larger than the intraspecific divergences and these patterns are stable over wide geographic ranges of Europe. In measureable terms, the genetic divergences between the species, based on Kimura 2-parameter (Kimura 1980) from Carolan et al. (2012) ranged from 0.033 to 0.044, whereas intraspecific distance was from 0.002 to 0.004. In the analysis by Murray et al. (2008), which was based on Tamura-Nei (Tamura and Nei 1993), they are slightly smaller. The interspecific distance ranges from 0.023-0.036 and intraspecific from 0.001-0.004. Based on their divergences and missing intermediates, Murray et al. (2008) concluded that their results “provide strong support for the existence of *B. cryptarum*, *B. lucorum*, *B. magnus* and *B. terrestris* as species that are discrete genotypic clusters” with respect to the Genotypic Cluster Concept of species (Mallet 1995).

Additionally, the COI sequences are suitable for inexpensive and fast analyses with restriction fragment length polymorphisms (RFLP), if only the species identity and not the individual sequence is of interest. Therefore Murray et al. (2008) provided a protocol which was successfully applied in Waters et al. (2011). An enhanced version was published recently (Vesterlund et al. 2014). This more time consuming approach works well with smaller COI fragments and hence is better suited for degraded DNA. However, it should be mentioned that due to different primers, none of the RFLPs protocols worked with the so-called *Folmer region* (derived from the primers presented in Folmer et al. (1994)), which is widely used for DNA ‘barcode’ collections such as BOLD (Ratnasingham and Hebert 2007).

In conclusion, both the labial gland secretion profiles and the results from the analyses of the nucleotide sequences reveal sufficient support for three distinct species in the European *B. lucorum*-complex. Additional indications arise from the morphological and phenological implications and the cross-breeding experiments. To further enhance our knowledge in this respect, the investigation of nuclear genes of the three species is urgently needed and will confidently be a key issue in understanding the closer phylogenetic relationships in the species complex. In the best case, data from nuclear genes may help clarify the status of the described subspecies of *B. cryptarum* (cf. Rasmont 1984).

## Can the species be distinguished by morphology?

While the biochemical and genetic methods for determination are widely accepted today, the published information on the morphological distinguishability is confusing. Fortunately the combination of the methods used nowadays allow for the verification or invalidation of potential traits. Currently, the key in Rasmont (1984) is the most important reference for the determination of females since most other keys (e.g. Mauss 1994, Amiet 1996, Bertsch et al. 2004, Dorow 2004) share crucial traits with that of Rasmont or are based on it. In general, the characters of coloration have been examined much more intensively. It should be mentioned that in using Rasmont (1984), the entirety of characters are only cognizable in queens. In this respect, the occurrence of the first collar is particularly important, since this may be the only character that is accessible in the field (Rasmont 1984, Bertsch 1997, Bertsch et al. 2004).

### Identification of queens

Three distinct forms of the first collar have been suggested to identify queens from the *B. lucorum*-complex. The first describes the lateral border of the yellow collar, which has been mentioned as a characteristic trait many times (e.g. Skorikov 1913, Ball 1914, Trautmann and Trautmann 1915, Alford 1975, Rasmont 1981b, Rasmont 1984, Amiet 1996, Bertsch 1997, Bertsch et al. 2004). If the border extends down onto the episternum, it is associated with *B. magnus* (Fig. 1c) and *B. cryptarum* (Fig. 1b). For *B. magnus*, the collar was reported to extend far below the tegulae and become very broad below them. In contrast, a higher lateral border that is almost exclusively restricted to the pronatallobus points to *B. lucorum* (Fig. 1a & 1d). In the literature, this trait is often vaguely described as “below tegula” or not, which is not entirely correct, since the border of the episternum is slightly below the tegula (Fig. 1d). The second trait is a so-called “S” or “5” shape that can be found within the collar. The pile along the border of the pronatallobus and the episternum is black and forms the “S” (Fig. 1b). It is associated with *B. cryptarum*. Another hint comes from a strong melanization of the collar which has been reported for *B. cryptarum*. However, this trait is regionally restricted (Bertsch et al. 2004, Bertsch et al. 2005) and on rare occasions can occur in the other species, too (Carolan et al. 2012).

**Figure 1:**
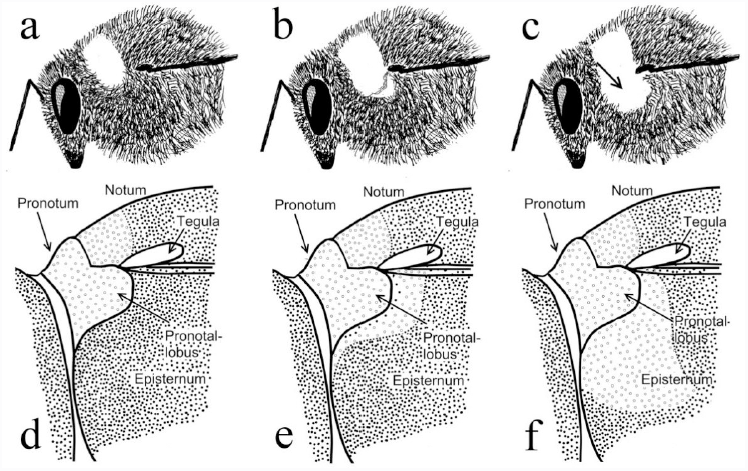
The shape of the first collar as described in the literature: *B. lucorum* (a, d), *B. cryptarum* (b, e) and *B. magnus* (c, f). Drawings from Neumayer (unpublished), based on Bertsch et al. (2004).

Bertsch (2009) was able to assign all but three investigated queen specimens to the correct species with the above mentioned characters, according to the biochemical evidence (n = 28). In contrast, using a larger sample from the British Isles and Denmark (n = 67), Carolan et al. (2012) showed that especially the collar-characters are not reliable for species diagnosis since they show overlap (see Fig. 4: doi:10.1371/journal.pone.0029251.g004). However, not every voucher of this figure is convincing: A close look at specimen “h” from their study, identified by morphology as “*lucorum*”, reveals an obvious collar extension far below the tegula and onto the episternum. There is no “S”-shape, it should therefore be associated with *B. magnus*, which is actually the case according to the DNA barcode. Moreover, specimen “c”, which was identified as “*magnus*” based on morphology reveals a faint black “S”-shape, exactly as described in Bertsch et al. (2004). It remains unclear why this voucher was assigned to *B. magnus* and not *B. cryptarum.* Thirdly, specimen “f” is not a typical *B. magnus*-morphotype since it does not show a clear broad collar below the tegula. Against this background, their conclusion that “each species can be morphologically identified as belonging to all 3 taxa” cannot be upheld. The study gave sufficient evidence that the extension of the anterior band of *B. cryptarum* queens can vary and that it critically resembles the traits of the other species. Yet it does not show that queens of *B. lucorum* and *B. magnus* resemble each other.

Aside from this confusing information, the work of Carolan et al. (2012) strongly indicated that the collar characters are not completely reliable and should not be exclusively taken into account for species identification. In addition, the key of Rasmont (1984) uses several characters aside from the coloration of the pile, such as the form of the labrum, punctuations of several structures and the shape of the hindleg metatarsus. However, the reliability of these characters has not been examined against a biochemical control and on an adequate scale. Thus the distinguishability of queens of the *B. lucorum*-complex can solely be presumed, even if most specimens are probably easily determined as described previously (Bertsch 1997, Bertsch et al. 2004, Bertsch et al. 2005).

### Identification of workers

The current state of knowledge concerning the identification of workers is worse than that for queens. Rasmont (1984) postulated that the coloration of workers corresponds approximately to the coloration of queens, implying a potential distinguishability in the shape and extension of the first collar. Unfortunately, the “S”-shape of *B. cryptarum* workers can be inconspicuous or absent (P. Rasmont, personal communication). Indications for the recognition of *B. magnus* can arise if yellow hair is mixed in the black pile of the first tergum (Rasmont 1984). Recently, the distinguishability of the anterior yellow band was examined quantitatively with Scottish (Waters et al. 2011) and Austrian specimens (Bardakji 2013) and was verified with RFLPs and DNA barcodes, respectively. Both studies revealed an uncertain connection of the traits to the species. In Scotland, where all three species occur sympatrically, Waters et al. (2011) was unable to properly distinguish *B. cryptarum* from the other two species. Still there was significant difference in collar extension between *B. lucorum* and *B. magnus*. It seems that the collar extension of the Scottish *B. cryptarum* is moderately variable and therefore constrains the possibility to recognize the other two species. Unfortunately, data on the pile coloration of the first tergum were not recorded, therefore the accuracy of this potential character remains uncertain.

In a study with Austrian specimens (Bardakji 2013), the sample consisted of *B. lucorum* and *B. cryptarum* individuals only. Regarding the extension of the collar, Bardakji (2013) was able to identify a great part (85.5%, 47 of n = 55) of the workers correctly. There were considerably more identification errors in *B. cryptarum*, supporting the view that the extension of the collar of workers of *B. cryptarum* is more variable in contrast to the others species. Aside from coloration, Rasmont (1984) described two groups of morphological characters that are accessible in queens and workers. (**I**) The first distinguishes characters of the labrum, e. g., the form of the basal area, especially if it is “U”-shaped (*B. lucorum* and *B. magnus*, Fig. 2a and 2c, respectively) or “V”-shaped (*B. cryptarum*, Fig. 2b). Further, the form of the lamella and punctuation are additional characters of potential value. (**II**) The second group describes the punctuations of the second tergum. Based on this, it was possible only to distinguish *B. lucorum* but not *B. cryptarum* or *B. magnus*. In contrast, Bardakji (2013) tested the reliability of the tergum-trait to differentiate between *B. lucorum* and *B. cryptarum.* It failed in roughly 1 of 5 cases. This is in line with the view of Dorow (2004), who challenged this character by describing a greater variation of the second tergum than previously described (Rasmont 1984). In any case, as mentioned above, *B. magnus* was not present in the sample used by Bardakji (2013) and therefore no general statements can be made. Still, it is strongly advised to test these traits on a larger scale with all three species. To avoid misunderstandings it is important to separately name the essential structures. The lamella is the structure directly below the basal area of the labrum and is neither “U” nor “V”-shaped. These shapes refer instead to the basal elevation of the labrum (Fig. 2).

**Figure 2:**
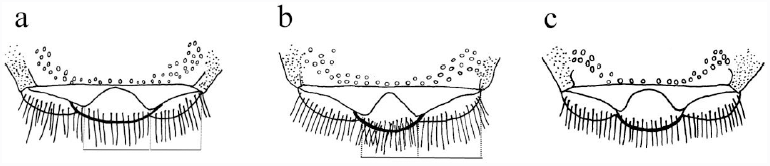
Labrum characters of *B. lucorum* (a), *B. cryptarum* (b) and *B. magnus* (c). Drawings taken from Neumayer (unpublished), based on Rasmont (1984).

In summary, the possibility to identify workers of all three species based on morphology has not been verified. Nonetheless, the characters of the labrum and the second tergum are particularly promising. Further morphological comparative examinations, which are supported by DNA barcoding, are necessary to verify these potential identifying characters and to uncover new traits. In the field, the extension of the first collar may be an indicator but is definitely not reliable, especially if all three species co-occur. Additionally, the reliability of the yellowish coloration of the first tergum for workers of *B. magnus* is worthy of further investigation.

### Identification of males

Identification of the males is probably the most difficult caste. Authors of recently published studies agreed that they are indistinguishable by morphology (Murray et al. 2008, Bertsch 2009, Waters et al. 2011). All three taxa are very similar and show extensive and overlapping variation in color and male genitalia (Rasmont et al. 1986). Therefore, keys based on coloration of the pile of the “face” (e.g. Amiet 1996, Dorow 2004, Gokcezade et al. 2010) are of restricted value, even if they may work for certain geographic regions. In the wider European context, these keys will fail to properly distinguish all male specimens of the complex. Aggravating this situation is the fact that the males of *B. terrestris* may also be confused with males of the *B. lucorum*-complex, in particular with specimens of *B. cryptarum* that have a dark facial pile. Although, *B. cryptarum* males often show the “S”-shape, it is geographically restricted and especially *B. cryptarum* and *B. magnus* can be more or less identical in morphology (P. Rasmont, personal communication). Aside from coloration, Rasmont et al. (1986) described several potential morphological characters. In this respect, the authors highlighted the punctuation of the second tergum as a distinguishing character for *B. lucorum* against *B. cryptarum* and *B. magnus*. Additional characters concern the diameter of the ocelli and the shape of the eighth tergum. The reliability of these traits in the wider European context remains uncertain. As long as no new insights in the distinguishability of the males are gained, completely reliable identification can only be achieved by biochemical or genetic approaches.

## Current challenges

### Difficulty in species recognition constrains our current knowledge

The long and difficult process of the recognition and acceptance of the species of the *B. lucorum*-complex has caused a number of critical problems concerning our current knowledge of the species ecology and distribution. First, the lack of applicable characters that are useful and easy for identification makes it difficult to obtain reliable data from the literature. The great majority of previous studies on these species are based on morphological identification and hence should be considered critically. Additionally, the possibilities of achieving faunistic data by interested amateurs and citizen scientists are very restricted and can barely contribute to scientifically founded statements in this concern. Second, the late redescription of *B. cryptarum* by Rasmont (1981a) implies that practically all data published before the redescription is unreliable since it was not possible to distinguish the species based on the debatable morphological traits. An example from Austria illustrates this point. All reported findings of *B. magnus* from Austria known to the author either before or shortly after the redescription, including the records of Schedl (1982) and Mathis (1982), findings from W. F. Reinig in Aistleitner (2000) and Ressl (1995), and the findings from B. Tkalců in Neumayer and Kofler (2005), were recently reexamined and found to belong to *B. cryptarum* based on morphology (J. Neumayer, personal communication). This demonstrates the importance of verifying older records from the literature and reveals that the uncommented use of references published before that date would lead to confusion, such as the citation of the textbook of Alford (1975) in Murray et al. (2008) or Waters et al. (2011). Third, the predicament is additionally aggravated by the individual treatment of *B. cryptarum* by the respective authors. Several experts declined to immediately accept *B. cryptarum* as a valid taxon and pooled the available data. In a strict sense, the identification method of every contribution should be reexamined, and the information in several reference textbooks or compilations (e.g. Prŷs-Jones and Corbet 1987, Westrich 1990) unfortunately cannot be regarded as reliable. In light of these problems, the number of applicable studies is strongly reduced. Reliable ecological and distributional data is primarily available in recent studies based on biochemical identification methods. Further, the excellent publications of Pierre Rasmont (Rasmont 1981a, Rasmont 1983, Rasmont 1984, Rasmont et al. 1986, Banaszak and Rasmont 1995) deserve our continued attention regarding the bumblebees of the *B. lucorum*-complex.

### Current issues concerning the distribution and ecology

Bertsch et al. (2004) carefully outlined the distribution of the species. Additional data comes from the recent COI-based studies (Murray et al. 2008, Anagnostopoulos 2009, Bertsch 2009, Waters et al. 2011, Carolan et al. 2012, Vesterlund et al. 2014) and from the distribution maps on bumblebees in the Atlas Hymenoptera project (Rasmont and Pauly 2010, Rasmont and Iserbyt 2010-2013). In these works the distribution, especially of *B. cryptarum* and *B. magnus*, appears fragmentary. The isolated finding of *B. cryptarum* in the Balkans (Anagnostopoulos 2009), the lack of doubtless identified *B. magnus* from the Iberian Peninsula south of the Pyrenees and the old records from Eastern Europe reveals the need of further sampling in these regions. Specimens from the Iberian Peninsula are of particular importance since there are indications that queens of *B. lucorum* exhibit a collar coloration similar to *B. magnus* queens in central Spain (Bertsch 2009). Against the background of the false records from the Austrian Alps, the presence of *B. magnus* in the Western Alps and along the southern slopes of the Alps must be verified as well. Species identification accomplished using COI barcodes should contribute to clarify our patchy knowledge on the distribution of the genetic haplotypes and might help outline postglacial recolonization events.

Further investigations are also necessary to understand the factors that drive the species abundances on regional and European scales, since the species composition can vary greatly at the regional level (Murray et al. 2008, Waters et al. 2011). One factor that has been claimed to influence the species composition is altitude. For example, Murray et al. (2008) revealed a changing species composition along a relatively low altitudinal gradient, and Neumayer and Paulus (1999) regarded *B. cryptarum* to be a high mountain species. Further, Scholl and Obrecht (1983) concluded that one *B. lucorum* s.l. taxon occurs preferably in the Alps. In contrast, the fact that all species can be found to live sympatrically in various lowland habitats in greater parts of Europe means that altitude cannot be the determining factor for the species distribution. However, the association of *B. cryptarum* with the high altitudes of the Alps and the observation of Pamilo et al. (1997) that *B. cryptarum*/*B. magnus* becomes predominant in northernmost Finland gives a reasonable background to study decisive factors that change with increasing altitude and latitude. Additional research is still pending concerning the habitat usage and nesting biology. Waters et al. (2011) made significant inroads to understand habitat usage but the study was restricted to relatively few habitats in northwestern Scotland. Regarding continental Europe, most recent studies specify the habitat types of the collection areas, but comparative studies concerning the used habitat or nesting sites over a sufficient geographic area are not available. Especially the exact habitat usage of *B. magnus* appears questionable. The occurrence of this species seems to be very patchy but regionally common (Bertsch et al. 2004). Further, it is frequently associated with heathlands (Banaszak and Rasmont 1994, Waters et al. 2011) and visits species of Ericaceae (Rasmont 1984, Bertsch et al. 2004). In contrast, the species is neither restricted to heathland nor does it rely on Ericaceae. Of particular note is that the species seems to occur in habitats with a very low diversity of flowering plant species, such as mass-flowering Ericaceae in heathlands or *Melampyrum pratense* in commercial forests (personal observation). Comparative studies are also necessary to improve our knowledge of the species bionomies, e.g. by examining exact nesting sites, and might confirm the phenological differences supposed by Bertsch et al. (2004).

### The importance of regionally stable characters

Traditionally, a significant part of the faunistic data of bumblebees in Europe is contributed by dedicated amateurs from the public rather than institutional scientists. At present, the restrictive possibility of identifying specimens by morphology has prevented reports of species of the *Bombus lucorum*-complex by citizen science. However, observations described in the literature suggest that species of the complex exhibit certain characters in certain regions such as the characteristic melanized collar, probably restricted to queens of *B. cryptarum* in northern Germany (Bertsch et al. 2005), or the “pinkish-buff” on the metasoma of fresh *B. magnus* queens that was reported from Northern Scotland (Macdonald 1999). It should be worthwhile to consider the reliability and stability of such characters to allow the public to make use of them for morphologically based identifications in particular regions. In this respect, particularly promising is the coloration of clypeal hairs in males. Admittedly males show extensive color variation in facial hair (Rasmont et al. 1986, Table 2), yet there are indications that regionally stable characters exist. The keys of Amiet (1996), Dorow (2004) and Gokcezade et al. (2010) share the same color-based system to distinguish males of all species from Switzerland, Hessia (Germany) and Austria, respectively. An examination of the reliability of these traits in the mentioned regions is of particular value, since the relevant characters are accessible in the field and hence might serve as a window to achieve distributional data without the need of biochemical controls.

## Future tasks

1. Investigations of nuclear genes of the species from the *Bombus lucorum*-complex will contribute to underpin the species’ status and help to estimate more accurate phylogenies.
2. New genetic sequence data, especially from the Mediterranean peninsulas, will enhance the current knowledge about the genetic diversity within the complex and might help to evaluate potential postglacial recolonization events.
3. The reported distinguishing morphological characters must be tested for all castes of all species in a wider European context against a biochemical control and on a sufficiently large scale. Further, it would be of particular importance to discover new distinguishing characters.
4. Investigations to determine regionally stable morphological or coloration characters might facilitate the acquisition of new distributional and ecological data by citizen scientists.
5. The reexamination of museum specimens, at best, backed up with an estimation of COI fragments, can allow the correct assignment of historic records and will help to identify incorrect species identifications.
6. Additional acquisition of safe ecological and distributional data will increase our knowledge about the species’ ecology. Among others, future studies should focus on altitudinal differences, nesting sites and habitat usage of the species.

## Acknowledgements

I am deeply grateful to J. Neumayer for the constant provision of meaningful information and the valuable drawings. Further, I would like to wholeheartedly thank B.-A. Gereben-Krenn, H. W. Krenn and S. Bardakji for their great support and for making inroads into the topic. I am also greatly indebted to M. A. Macdonald and P. Rasmont for sharing useful information. Lastly, great thanks go to J. Plant for very helpful suggestions to the text.

## References

Aistleitner E (2000) Fragmenta entomofaunistica IV Daten zur Hautflügler-Fauna Vorarlbergs, Austria occ. (Insecta, Hymenoptera). Entomofauna 21 (19): 237–248.

Alford DV (1975) Bumblebees. Davis-Poynter (London): 1–352.

Amiet F (1996) Hymenoptera Apidae, 1. Teil - Allgemeiner Teil, Gattungsschlüssel, die Gattungen Apis, Bombus und Psithyrus. Insecta Helvetica Fauna (Neuchâtel, Switzerland): 1–98.

Amiet F, Krebs A (2012) Bienen Mitteleuropas - Gattungen, Lebensweise, Beobachtung. Haupt Verlag (Bern, Stuttgart, Wien): 1–424.

Anagnostopoulos IT (2009) New Balkan records of Bombus subterraneus (Linnaeus 1758) and Bombus cryptarum (Fabricius 1775) from Greece. Entomologia Hellenica 18: 56–61.

Ander K (1965) Über die Verbreitung der Hummeln in Schweden (Hym. Apidae). Opuscula Entomologica 30: 135–139.

Baker DB (1996) On a collection of Bombus and Psithyrus principally from Sutherland, with notes on the nomenclature or status of three species (Hymenoptera, Apoidea). British Journal of Entomology and Natural History 9: 7–19.

Ball JF (1914) Les bourdons de la Belgique. Annales de la Société entomologique de Belgique 58: 77–108.

Banaszak J, Rasmont P (1994) Occurrence and distribution of the subgenus Bombus Latreille sensu stricto in Poland (Hymenoptera, Apoidea). Polskie Pismo Entomologiczne 63: 337–356.

Bardakji S (2013) Identification of cryptic species belonging to the Bombus lucorum - complex: DNA barcoding and morphological approaches. Master’s Thesis at the University of Vienna (Austria).

Bergström G, Kullenberg B, Ställberg-Stenhagen S (1973) Studies on natural odoriferous compounds. VII. Recognition of two forms of Bombus lucorum L. (Hymenoptera, Apidae) by analysis of the volatile marking secretion from individual males. Chemica Scripta 4: 174–182.

Bergström G, Svensson BG, Appelgren M, Groth I (1981) Complexity of Bumble Bee Marking Pheromones: Biochemical, Ecological and Systematical Interpretations. In: Howse PE, Clément J-L (Eds.) Biosystematics of Social Insects. Systematics Association Special Volume No. 19, Academic Press (London and New York): 175–183.

Bertsch A (1997) Abgrenzung der Hummel-Arten Bombus cryptarum und B lucorum mittels männlicher Labialdrüsen-Sekrete und morphologischer Merkmale (Hymenoptera, Apidae). Entomologia Generalis 22 (2): 129–145.

Bertsch A (2009) Barcoding cryptic bumblebee taxa: B. lucorum, B. crytarum and B. magnus, a case study (Hymenoptera: Apidae: Bombus). Beiträge zur Entomologie 59 (2): 287–310.

Bertsch A, Schweer H (2012) Male labial gland secretions as species recognition signals in species of Bombus. Biochemical Systematics and Ecology 40: 103–111. doi: 10.1016/j.bse.2011.10.009.

Bertsch A, Schweer H, Titze A (2004) Discrimination of the bumblebee species Bombus lucorum, B. cryptarum and B. magnus by morphological characters and male labial gland secretions (Hymenoptera: Apidae). Beiträge zur Entomologie 54 (2): 365–386.

Bertsch A, Schweer H, Titze A, Tanaka H (2005) Male labial gland secretions and mitochondrial DNA markers support species status of Bombus cryptarum and B. magnus (Hymenoptera, Apidae). Insectes Sociaux 52 (1): 45–54.

Bučánková A, Komzáková O, Cholastová T, Ptáček V (2011) Notes on distribution of Bombus cryptarum (Hymenoptera, Apoidea) in Moravian territory (Czech Republic) and its laboratory rearing. Acta Universitatis Agriculturae et Silviculturae Mendelianae Brunensis 59 (6): 69–74.

Carolan JC, Murray TE, Fitzpatrick Ú, Crossley J, Schmidt H, Cederberg B, McNally L, Paxton RJ, Williams PH, Brown MJF (2012) Colour Patterns Do Not Diagnose Species: Quantitative Evaluation of a DNA Barcoded Cryptic Bumblebee Complex. PLoS ONE 7 (1): e29251. doi: 10.1371/journal.pone.0029251.

Corbet SA, Williams IH, Osborne JL (1991) Bees and the pollination of crops and wild flowers in the European Community. Bee World 72 (2): 47–59.

De Jonghe R (1982) Copulations interspécifiques en captivité d’especes du genre Bombus Latreille (sensu stricto)(Hymenoptera, Apidae, Bombinae). Bulletin et Annales de la Société Royale Belge d’Entomologie 118: 171–175.

De Jonghe R, Rasmont P (1983) Kreuzungsexperiment mit Hummeln des Genus Bombus Latreille sensu stricto (Hymenoptera, Apidae). Phegea 11 (1): 7–10.

Delmas R (1981) Systematics and Geographical Variation in the Bombinae. In: Howse PE, Clément J-L (Eds.) Biosystematics of Social Insects. Systematics Association Special Volume No. 19, Academic Press (London and New York): 223–229.

DeSalle R, Egan MG, Siddall M (2005) The unholy trinity: taxonomy, species delimitation and DNA barcoding. Philosophical Transactions of the Royal Society. Series B: Biological Sciences 360 (1462): 1905–1916. doi: 10.1098/rstb.2005.172.

Dorow WHO (2004) Hymenoptera: Aculeata (Stechimmen). In: Dorow WHO, Flechtner G, Kopelke JP (Eds.) Naturwaldreservate in Hessen 6/2.2. Schönbuche. Zoologische Untersuchungen 1990- 1992, Teil 2. Hessen-Forst - Forsteinrichtung, Information, Versuchswesen, Giessen & Forschungsinstitut Senckenberg. FIV Ergebnis- und Forschungsbericht (Frankfurt, Germany): 127–264.

Elfving R (1960) Die Hummeln und Schmarotzerhummeln Finnlands. Fauna Fennica 10: 1–43.

Folmer O, Black M, Hoeh W, Lutz R, Vrijenhoek R (1994) DNA primers for amplification of mitochondrial cytochrome c oxidase subunit I from diverse metazoan invertebrates. Molecular Marine Biology and Biotechnology 3 (5): 294–299.

Gokcezade JF, Gereben-Krenn B-A, Neumayer J, Krenn HW (2010) Feldbestimmungsschlüssel für die Hummeln Österreichs, Deutschlands und der Schweiz (Hymenoptera: Apidae). Linzer biologische Beiträge 42 (1): 5–42.

Goulson D (2010) Bumblebees: behaviour, ecology, and conservation. Oxford University Press (New York): 1–317.

Goulson D, Lye GC, Darvill B (2007) Decline and Conservation of Bumble Bees. Annual Review of Entomology 53: 191–208. doi: 10.1146/annurev.ento.53.103106.093454.

Kimura M (1980) A Simple Method for Estimating Evolutionary Rates of Base Substitutions Through Comparative Studies of Nucleotide Sequences. Journal of Molecular Evolution 16: 111–120.

Krüger E (1951) Phänoanalytische Studien an einigen Arten der Untergattung Terrestribombus O. Vogt (Hymen. Bomb.). I. Teil. Tijdschrift voor Entomologie 93: 141–197.

Krüger E (1954) Phaenoanalytische Studien an einigen Arten der Untergattung Terrestribombus O. Vogt (Hymenoptera, Bombidae). II. Teil. Tijdschrift voor Entomologie 97: 263–298.

Kullenberg B, Bergström G, Ställberg-Stenhagen S (1970) Volatile Components of the Cephalic Marking Secretion of Male Bumble Bees. Acta Chemica Scandinavica 24 (4): 1481–1483.

Løken A (1973) Studies on Scandinavian Bumble Bees (Hymenoptera, Apidae). Norsk Entomologisk Tidsskrift 20 (1): 1–219.

Macdonald MA (1999) A contribution to the Bombus magnus/lucorum debate. BWARS Newsletter Autumn 1999 (2): 9.

Mallet J (1995) A species definition for the Modern Synthesis. Trends in Ecology & Evolution 10 (7): 294–299.

Mathis H (1982) Untersuchungen zur Phänologie und Arealkunde heimischer Arten der Gattung Bombus (Hummeln). Lehramtshausarbeit. Pädagogischen Hochschule Vorarlberg (Feldkirch).

Mauss V (1994) Bestimmungsschlüssel für Hummeln. Deutscher Jugendbund für Naturbeobachtung (Hamburg): 1–50.

Meyer CP, Paulay G (2005) DNA Barcoding: Error Rates Based on Comprehensive Sampling. PLoS Biology 3 (12): e422. doi: 10.1371/journal.pbio.0030422.

Michener CD (2007) The Bees of the World. The Johns Hopkins University Press (Baltimore): 1–953.

Murray TE, Fitzpatrick U, Brown MJF, Paxton RJ (2008) Cryptic species diversity in a widespread bumble bee complex revealed using mitochondrial DNA RFLPs. Conservation Genetics 9: 653–666. doi: 10.1007/s10592-007-9394-z.

Neumayer J, Kofler A (2005) Zur Hummelfauna des Bezirkes Lienz (Osttirol, Österreich)(Hymenoptera: Apidae, Bombus). Linzer biologische Beiträge 37 (1): 671–699.

Neumayer J, Paulus HF (1999) Ökologie alpiner Hummelgemeinschaften: Blütenbesuch, Ressourcenaufteilung und Energiehaushalt. Untersuchungen in den Ostalpen Österreichs. Stapfia 67: 5–246.

Obrecht E, Scholl A (1984) Bombus lucorum auct. ein Artenkomplex - Enzymelektrophoretische Befunde (Hymenoptera, Bombidae). Verhandlungen der Deutschen Zoologischen Gesellschaft 77: 266.

Pamilo P, Tengö J, Rasmont P, Pirhonen K, Pekkarinen A, Kaarnama E (1997) Pheromonal and enzyme genetic characteristics of the Bombus lucorum species complex in northern Europe. Entomologica Fennica 7: 187–194.

Pamilo P, Varvio-Aho S-L, Pekkarinen A (1984) Genetic variation in bumblebees (Bombus, Psithyrus) and putative sibling species of Bombus lucorum. Hereditas 101: 245–251.

Pekkarinen A (1979) Morphometric, colour and enzyme variation in bumblebees (Hymenoptera, Apidae, Bombus) in Fennoscandia and Denmark. Acta Zoologica Fennica 158: 1–60.

Prŷs-Jones OE, Corbet SA (1987) Bumblebees. Cambridge University Press (Cambridge):1–86.

Rasmont P (1981a) Redescription d’une espèce méconnue de bourdon d’Europe: Bombus lucocryptarum Ball, 1914 N. Status (Hymenoptera, Apidae, Bombinae). Bulletin et Annales de la Société Royale Belge d’Entomologie 117: 151–154.

Rasmont P (1981b) Contribution à l’étude des bourdons du genre Bombus Latreille, 1802 sensu stricto (Hymenoptera, Apidae, Bombinæ). Travail de fin d’études. Faculté des Sciences Agronomiques de l’État Gembloux (Belgium).

Rasmont P (1983) Notes Taxonomiques sur les Bourdons. Bulletin et Annales de la Société royale belge d’Entomologie 119: 167–170.

Rasmont P (1984) Les bourdons du genre Bombus Latreille sensu stricto en Europe Occidentale et Centrale (*Hymenoptera, Apidae*). Spixiana 7 (2): 135–160.

Rasmont P, De Jonghe R (1985) Progrès récents dans la connaissance des bourdons du genre Bombus Latreille sensu stricto (Hymenoptera, Apidae, Bombinae). Actes des Colloques Insectes Sociaux 2: 119–122.

Rasmont P, Iserbyt S (2010-2013) Atlas of the European Bees: genus Bombus. 3rd Edition, Atlas Hymenoptera (Mons, Gemloux). http://www.zoologie.umh.ac.be//hymenoptera/page.asp?ID=169.

Rasmont P, Pauly A (2010) Les bourdons de la Belgique: Atlas Hymenoptera (Mons, Gembloux). http://www.zoologie.umh.ac.be//hymenoptera/page.asp?ID=160.

Rasmont P, Scholl A, De Jonghe R, Obrecht E, Adamski A (1986) Identité et variabilité des mâles de bourdons du genre Bombus Latreille sensu stricto en Europe occidentale et centrale (Hymenoptera, Apidae, Bombinae). Revue suisse de Zoologie 93 (3): 661–682.

Ratnasingham S, Hebert PDN (2007) BOLD: The Barcode of Life Data System (http://www.barcodinglife.org). Molecular Ecology Notes 7 (3): 355–364. doi: 10.1111/j.1471-8286.2006.01678.x.

Reinig WF (1939) Die Evolutionsmechanismen, erläutert an den Hummeln. Verhandlungen der Deutschen Zoologischen Gesellschaft Supplement 12: 170–206.

Ressl F (1995) Naturkunde des Bezirkes Scheibbs - Tierwelt (3). Scheibbs, Naturkundliche Arbeitsgemeinschaft (Linz/Österreich): 1–444.

Schedl W (1982) Über aculeate Hautflügler der zentralen Ötztaler Alpen (Tirol, Österreich) (Insecta: Hymenoptera). Berichte des naturwissenschaftlich-medizinischen Vereins in Innsbruck 69: 95–117.

Scholl A, Obrecht E (1983) Enzymelektrophoretische Untersuchungen zur Artabgrenzung im Bombus lucorum-Komplex (Apidae, Bombini). Apidologie 14 (2): 65–78.

Scholl A, Obrecht E, Owen RE (1990) The genetic relationship between Bombus moderatus Cresson and the Bombus lucorum auct. species complex (Hymenoptera: Apidae). Canadian Journal of Zoology 68 (11): 2264–2268.

Scholl A, Thorp RW, Obrecht E (1992) The genetic relationship between Bombus franklini (Frison) and other taxa of the subgenus Bombus s. str. (Hymenoptera: Apidae). Pan-Pacific Entomologist 68: 46–51.

Skorikov A (1913) Neue Hummelformen (Hymenoptera, Bombidae). Revue Russe d’Entomologie 13: 171–175.

Tamura K, Nei M (1993) Estimation of the Number of Nucleotide Substitutions in the Control Region of Mitochondrial DNA in Humans and Chimpanzees. Molecular Biology and Evolution 10 (3): 512–526.

Taylor HR, Harris WE (2012) An emergent science on the brink of irrelevance: a review of the past 8 years of DNA barcoding. Molecular Ecology Resources 12 (3): 377–388. doi: 10.1111/j.1755-0998.2012.03119.x.

Tkalců B (1974) Bemerkenswerte Bienenfunde in der Tschechoslowakei (Hymenoptera, Apoidea). Acta entomologica Bohemoslovaca 71: 205–208.

Trautmann G, Trautmann W (1915) Bombus terrestris L. var. nov. flavoscutellaris. Internationale Entomologische Zeitschrift 1915: 18.

Velthuis HHW, van Doorn A (2006) A century of advances in bumblebee domestication and the economic and environmental aspects of its commercialization for pollination. Apidologie 37 (4): 421–451. doi: 10.1051/apido:2006019.

Vesterlund S-R, Sorvari J, Vasemägi A (2014) Molecular identification of cryptic bumblebee species from degraded samples using PCR–RFLP approach. Molecular Ecology Resources 14 (1): 122–126. doi: 10.1111/1755-0998.12168.

Vogt O (1909) Studien über das Artproblem. 1. Mitteilung. Über das Variieren der Hummeln. 1. Teil. Sitzungsberichte der Gesellschaft naturforschender Freunde zu Berlin 1909: 28–84.

Vogt O (1911) Studien über das Artproblem. 2. Mitteilung. Über das Variieren der Hummeln. 2. Teil. (Schluss). Sitzungsberichte der Gesellschaft naturforschender Freunde zu Berlin 1911: 31–74.

von Dalla Torre KW (1880) Unsere hummel-(Bombus) Arten. Der Naturhistoriker 2: 40–41.

Warncke K (1981) Die Wildbienen des Klagenfurter Beckens (Hymenoptera, Apidae). Carinthia II 171 (91): 275–348.

Warncke K (1986) Die Wildbienen Mitteleuropas, ihre gültigen Namen und ihre Verbreitung (Insecta: Hymenoptera). Entomofauna Supplement 3: 1–128.

Waters J, Darvill B, Lye GC, Goulson D (2011) Niche differentiation of a cryptic bumblebee complex in the Western Isles of Scotland. Insect Conservation and Diversity 4: 46–52. doi: 10.1111/j.1752-4598.2010.00101.x.

Westrich P (1990) Die Wildbienen Baden-Württembergs I und II. - 2. verb. Auflage. Verlag Eugen Ulmer (Stuttgart): 1–972.

Will KW, Rubinoff D (2004) Myth of the molecule: DNA barcodes for species cannot replace morphology for identification and classification. Cladistics 20 (1): 47–55. doi: 10.1111/j.1096-0031.2003.00008.x.

Williams PH (1991) The bumble bees of the Kashmir Himalaya (Hymenoptera: Apidae, Bombini). Bulletin of the British Museum of Natural History (Entomology) 60 (1): 1–204.

Williams PH (1998) An annotated checklist of bumble bees with an analysis of patterns of description (Hymenoptera: Apidae, Bombini). Bulletin of the Natural History Museum London (Entomology) 67: 79–152.

Williams PH (2000) Are Bombus lucorum and magnus separate species? BWARS Newsletter Spring 2000 (1): 15–17.

Williams PH (2007) The distribution of bumblebee colour patterns worldwide: possible significance for thermoregulation, crypsis, and warning mimicry. Biological Journal of the Linnean Society 92: 97–118. doi: 10.1111/j.1095-8312.2007.00878.x.

Williams PH, Brown MJF, Carolan JC, An J, Goulson D, Aytekin AM, Best LR, Byvaltsev AM, Cederberg B, Dawson R, Huang J, Ito M, Monfared A, Raina RH, Schmid-Hempel P, Sheffield CS, Šima P, Xie Z (2012) Unveiling cryptic species of the bumblebee subgenus Bombus s. str. worldwide with COI barcodes (Hymenoptera: Apidae). Systematics and Biodiversity 10 (1): 21–56. doi: 10.1080/14772000.2012.664574.

Winter K, Adams L, Thorp R, Inouye D, Day L, Ascher J, Buchmann S (2006) Importation of Non-Native Bumble Bees into North America: Potential Consequences of Using Bombus terrestris and Other Non-Native Bumble Bees for Greenhouse Crop Pollination in Canada, Mexico, and the United States: 1–33.

